# Transcriptomic Implications of Toxic Effects of Nanoparticles on Metabolic Pathways of Liver Cells

**DOI:** 10.1101/2023.11.24.568576

**Authors:** Merve Erden Tüçer, Nazlican Tunç, Suat Tüçer, Urartu Özgür Şafak Şeker

## Abstract

While nanoparticles find applications in various fields such as cosmetics, drug delivery, and medicine, they also exhibit significant drawbacks. The adverse effects remain inadequately comprehended and necessitate further analysis. This study focuses on assessing the impacts of diverse nanoparticles on HepaRG cell spheroids prior to conducting RNA-Seq analysis, cell viability assays were performed to determine the concentrations of NPs that induce toxicity while maintaining 80% cell viability—a concentration sufficient to cause toxicity without total cell death. To mimic the cellular microenvironment, HepaRG cell spheroids were generated and treated with four distinct nanoparticles of varying sizes. Subsequently, these spheroids underwent RNA-seq analysis to identify specific genes responsive to nanoparticle-induced toxicity. Our findings demonstrate that exposure to nanoparticles substantially modifies gene transcription. Notably, 38 differentially expressed genes were shared across all four types of nanoparticles. These genes were linked to distinct categories of Gene Ontology (GO) terms, thereby clarifying their functions and metabolic pathways through GO and KEGG pathway analyses. Remarkably, processes such as apoptosis, sensitivity to metal ions, hypoxia, and oxidative stress consistently exhibited significant enrichment across all nanoparticle types.

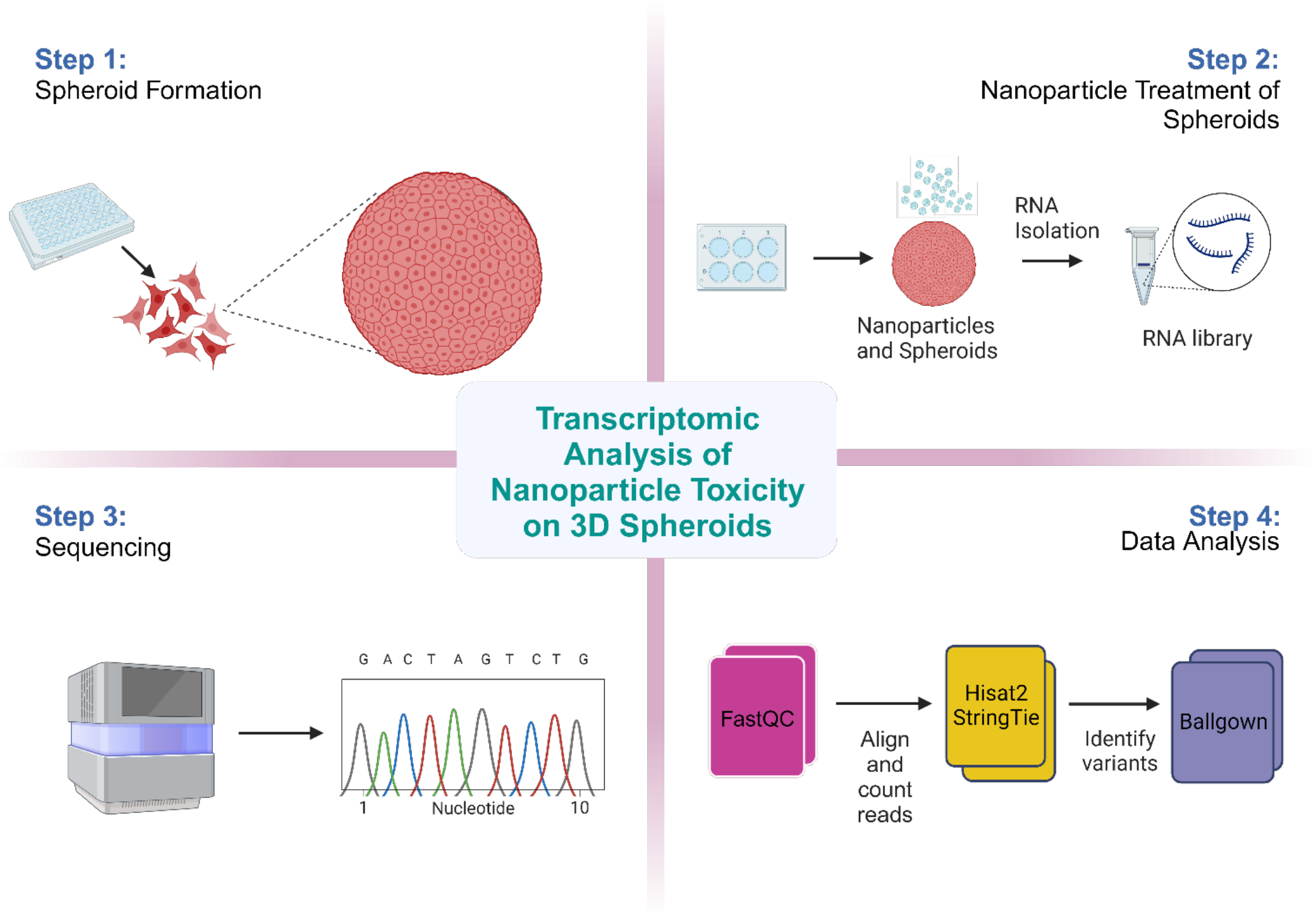

## Introduction

Nanotechnology has emerged as a prominent area of research, drawing attention since the last century, and at the forefront of this field are nanoparticles (NPs), the fundamental units of investigation. These materials generally constitute a vast category of matter up to 100 nm, differ in dimension including 0D, 1D, 2D, or 3D ^1, 2^. They can be made up of carbon, organic substances, metal or metal oxide and found in different shapes such as spherical, cylindrical, tubular, conical, hollow core, spiral, flat, or irregular structures ^3, 4^. NPs have unique properties due to large surface area, mechanical strength, optical activity, and chemical reactivity.^5^ These unique properties of NPs have led to their diverse applications in a wide array of fields, such as cosmetics, food and food packaging, drug delivery and medicine, bioremediation, paints, coatings, biosensing, and bioimaging.^6, 7^

Gold nanoparticles (NPs) have unique properties in terms of size and shape dependent optical features, and exhibit remarkable biocompatibility and chemical stability, making them highly sought after for a diverse range of applications in photonic device fabrication, catalysis, organic and biomolecules (bio)sensors, drug delivery, biological imaging, and therapy.^8,9,10^ Silver, renowned for its antimicrobial properties since ancient times, has found renewed relevance with the advancement of nanotechnology. Silver nanoparticles started to be exploited as an active biomedical factor as nanotechnology advanced.^11^ Thanks to their absorption and scattering features, they are using in biological sensing area as optical and spectroscopic tags. ^12^ In cancer treatments, titanium dioxide (TiO2) has been employed for its therapeutic potential. When TiO2 NPs are combined with targeted drugs, TiO2 induces DNA damage and apoptotic pathways. ^13^ Quantum dots (QDs), owing to their distinctive physical and optical characteristics and their ability to bind several biomolecules on their surface, serve as excellent candidates for biosensing applications and biological imaging.^14^ In the targeted drug delivery application, anticancer drugs specifically send to the tumor site while minimizing the potential harm to adjacent healthy cells.^15^ The utilization of magnetic nanoparticles (NPs) in drug delivery holds several advantageous features. For example, they can be visualized, and to induce drug delivery, they can be heated in magnetic field. ^16^

Although NPs offer numerous advantages, they also possess certain drawbacks. The high surface-to-volume ratio of NPs also makes them capable of causing cellular stress, which can result in gene expression disruption, protein unfolding, DNA damage, and reactive oxygen species (ROS) production, leading to various health issues.^18^ As human interactions with NPs continue to increase, comprehensive investigations into their impact on human health are essential. To assess the potential risk of injury to individuals exposed to nanoparticles, it is important to understand the intrinsic toxicity of the substance as well as the dose delivered to the target organ.^19^ Numerous studies have investigated the pro-inflammatory effects of nanoparticles as this property is a key indicator of their potential danger. One of the significant discoveries in this field is that nanoparticles have a more pronounced effect on inflammation, cell damage, and cell stimulation compared to an equivalent amount of larger-sized particles of the same substance.^20^ People are exposed to these commercially available nanoparticles through various everyday products. For example, clay nanoparticles can be found in beer, titanium dioxide (TiO2) nanoparticles are commonly used in sunscreens, carbon nanoparticles can be found in bicycles, and silver (Ag) nanoparticles are present in sheets and clothing.^21^ To gain an understanding of the potential risks associated with the use of nanoparticles in the food, food contact material and enzymes, an extensive literature search and inventory by the European Food Safety Authority (EFSA) resulted in the selection of 779 relevant references.^22^ These references are indication of negative effects of NPs for humans and environment. Also, the FDA has some criteria regarding the regulation of NPs. The evaluation of nanotechnology products by the Food and Drug Administration (FDA) takes into account any unique qualities and behaviors that may arise from their use. Two factors are considered when determining if FDA-regulated products incorporate nanotechnology. These factors include the size of the particles and any dimension-dependent characteristics or phenomena associated with nanoscale materials. If a product meets either of these criteria, both the industry and the FDA must assess whether any special qualities or behaviors related to nanotechnology have been identified and addressed during studies on the product’s safety, effectiveness, public health impact, or regulatory status. ^23^

In addition to their potential adverse impact on human health, nanoparticles also have detrimental effects on the environment. Naturally occurring nanoparticles in the environment originate from sources such as combustion products from burning fuels, atmospheric phytochemistry aerosols, and volcanic activity. ^1^ However, the impact of NPs on the environment is primarily determined by their ability to accumulate in organisms and to cause harm once they do. Unlike naturally occurring NPs, produced NPs contain surfactants and stabilizers that cause them to persist and not join together over time to form larger materials. Therefore, it is essential to assess the potential impact on the environment of using these materials. ^24^

The liver is a pivotal organ for investigating the toxicity of nanoparticles. While research on the subject has been extensive, the findings are somewhat constrained, necessitating a more intricate analysis of the underlying mechanisms. Therefore, a thorough pathway analysis is essential to enhance our comprehension of nanoparticle toxicity.

Although most hepatic cell lines have a limited capacity for bioactivation, they could serve as an alternative to primary hepatocytes. The HepaRG cell line, on the other hand, is the first human cell line capable of in vitro differentiation into mature hepatocyte-like cells.^25^ HepaRG cells are particularly attractive as a tool for investigating differentiation, liver metabolism, drug effect/metabolism/toxicity due to their progenitor nature, capacity to differentiate into biliary and hepatocyte phenotypes as well as the liver enzyme levels comparable to primary hepatocytes. ^26^ Tumors modelled in 2D cell cultures are overly simplistic, and they lack of tumor heterogeneity and microenvironment.^27^ The limitations inherent in two-dimensional (2D) systems have prompted the evolution toward three-dimensional (3D) in vitro systems, which provide in vivo-like settings and cell-cell and cell-extracellular matrix (ECM) interactions that are essential for the regulation of cell behaviour and function but are challenging to mimic in 2D. ^28^ 3D structures exhibit tissue-specific design, improved cell-cell and cell-extracellular matrix interactions, and responses to external stimuli that are representative of those seen in vivo.^29^ For the purpose of generating in vitro models that aptly replicate the intricacies of in vivo settings, the integration of three-dimensional (3D) structures has thus emerged as the critical component that has hitherto been absent.^30^ For these reasons, we choose to use spheroids rather than 2D culture.

RNA-seq is a combination of experimental techniques that uses library creation, massively parallel deep sequencing, and create cDNA sequences from the whole RNA molecules. Long non-coding RNA, miRNA, siRNA, and other small RNA classes (such as snRNA and piRNA) that are involved in regulating RNA stability, protein translation, or the manipulation of chromatin states have all been catalogued by RNA-seq.^31^ Gene expression, alternative splicing, and allele-specific expression are all better understood and quantified with RNA-Seq. Deep profiling of the transcriptome and the possibility to shed light on many physiological and pathological situations have been made possible by recent improvements in the RNA-Seq methodology, which span sample preparation, sequencing platforms, and bioinformatic data interpretation.^32^

In the present research, we employed four distinct types of nanoparticles (namely AuNP, AgNP, quantum dot (QD), and TiO2) in conjunction with HepaRG cells to establish a system to monitor nanoparticle cytotoxicity. By conducting transcriptome analysis, we obtain detailed information about how HepaRG cells reacted to the various nanoparticles’ cytotoxicity, as well as the genes whose expression was significantly changed.

## RESULTS

### Characterization of nanoparticles

Characterization of NPs in terms of their sizes is important for the proceeding analysis, because the size of the NPs alter their interaction with the cells and biomolecules. Not only the size of the nanoparticles, but the physicochemical properties of NPs alter their effect on the cells. To see the effects of the different sizes and materials on the nanoparticle properties on cells, we used 3 different materials of nanoparticles of 3 different sizes of each and these are gold (3nm, 15nm, 80nm), silver (12nm, 40nm, 80nm), titanium (100nm, 300nm and 500nm) and also we used 2 different quantum dots (CdZnSe/CdZnS/ZnS and InP/ZnS). The physicochemical properties of the NPs depend on the aggregation, dissolution, or coating with biological substances such as proteins. That is why the characterization of the NPs are done in the cell culture medium. UV-VIS emission spectra revealed the lambda max for each NPs. Peak absorbance for 3nm, 15nm and 80nm AuNPs were at 510nm, 520nm, and 550nm respectively (**Figure 1A**). For 12nm, 40nm and 80nm AgNPs the peak absorbance was at 400nm, 450nm, and 490nm respectively (**Figure 1B**). Also, UV-VIS emission spectra of each type and size of nanoparticle showed the size distribution of nanoparticles. The size and morphology of the nanoparticles were also characterized by transmission electron microscopy (TEM) (**Figure 1**).

**Figure 1.**
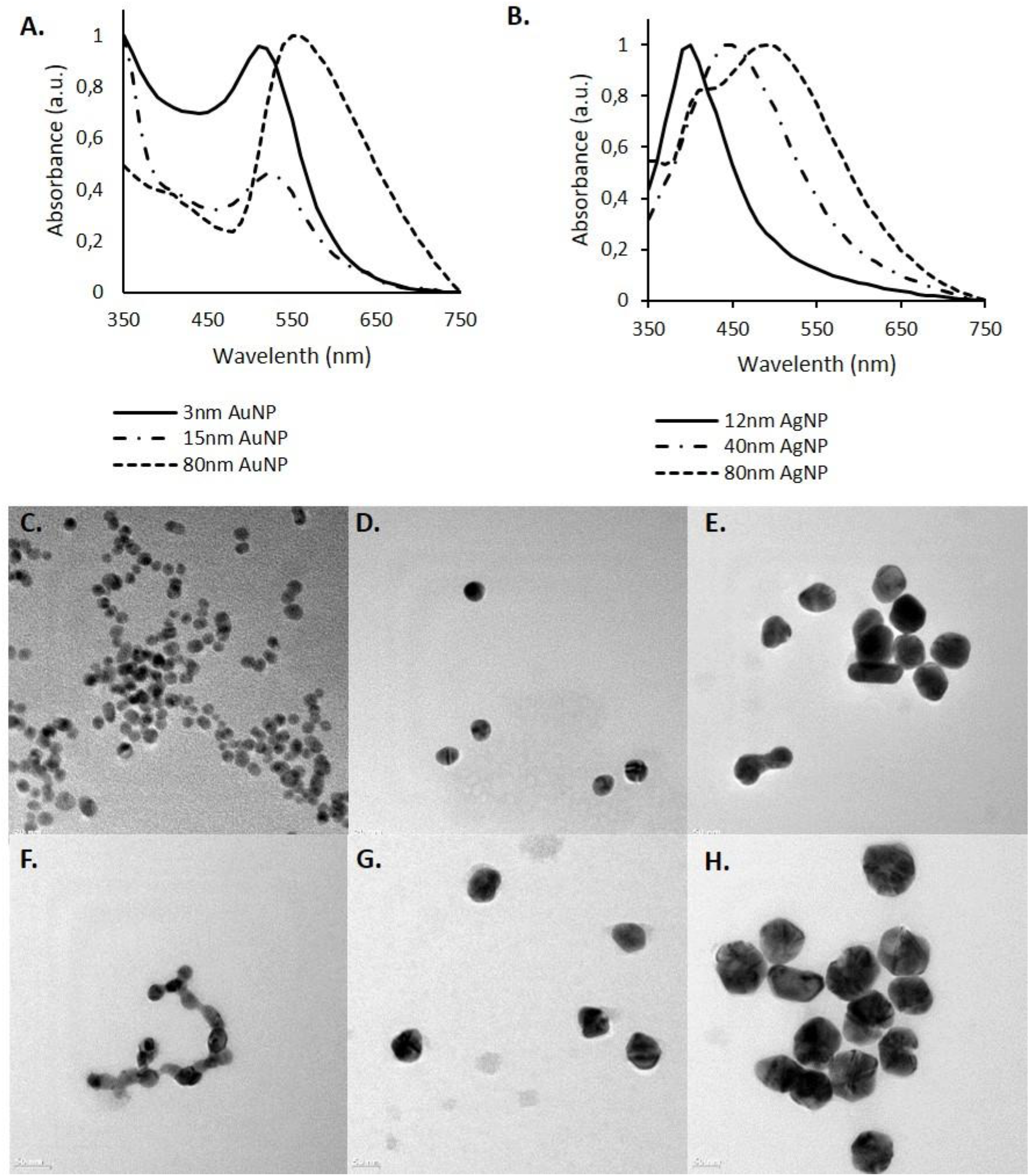
Characterization of NPs. UV-VIS emission spectra of **A)** AuNPs **B)** AgNPs. TEM images of nanoparticles **C)** 3nm AuNP **D)** 15nm AuNP **E)** 80nm AuNP **F)** 12nm AgNP **G)** 40nm AgNP **H)** 80nm AgNP

### Cell Viability Assays

In order to do the differential gene expression analysis for the cytotoxicity of the NPs, we need to expose the cells with toxic concentrations of NPs before RNA isolation. In this regard, we did cell viability assays to find the toxic concentrations of NPs of different materials and sizes on HepaRG cells. HepaRG cells were exposed to different doses of NPs for 24 hours, then the toxicity of the NPs was measured by MTT assay method. As we expected, cell viability decreased with increasing doses of NPs (**Figure 2**). We decided to choose the concentration as toxic concentration of NPs which we collect mRNAs at is to be the concentration at which the viability is 80%. We decided on this number, because we wanted to NP concentration to be toxic, however not too toxic to kill all the cells since we were going to collect mRNAs from these cells after NP exposure. Then we proceeded to collect the mRNAs after finding the toxic concentrations for each type and size of the NPs.

**Figure 2.**
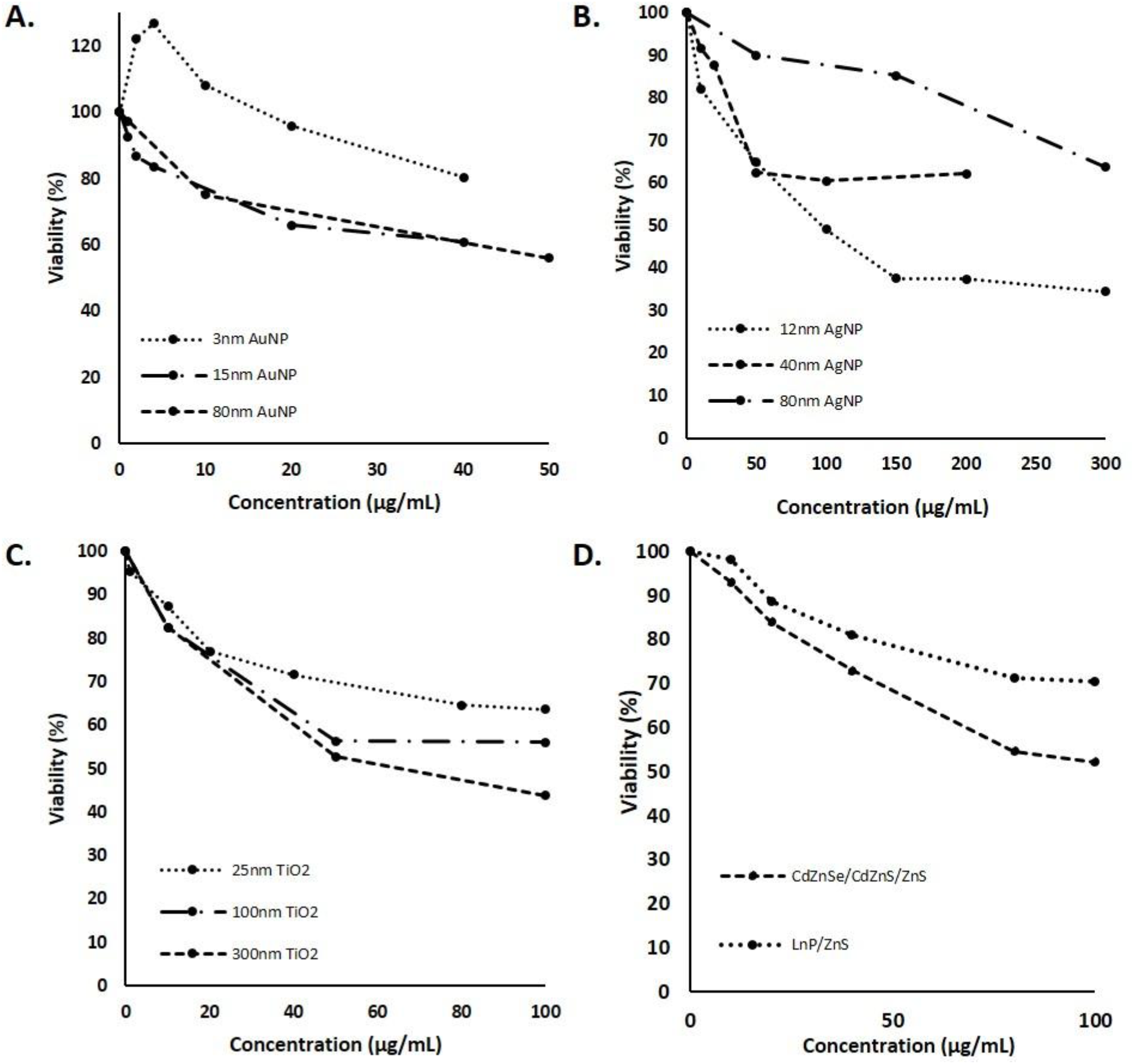
Cytotoxicity of nanoparticles on HepaRG cells for 24 hours. Cytotoxicity of **A)** Gold nanoparticles (3nm, 15nm, and 80nm) and **B)** Silver Nanoparticles (12nm, 40nm, 80nm) C) Titanium Nanoparticles (25nm, 100nm and 300nm) and D) QDs (InP/ZnS and CdZnSe/CdZnS/ZnS). Each value represents the mean ± SE of 3 repeats (N=3).

**Figure 3.**
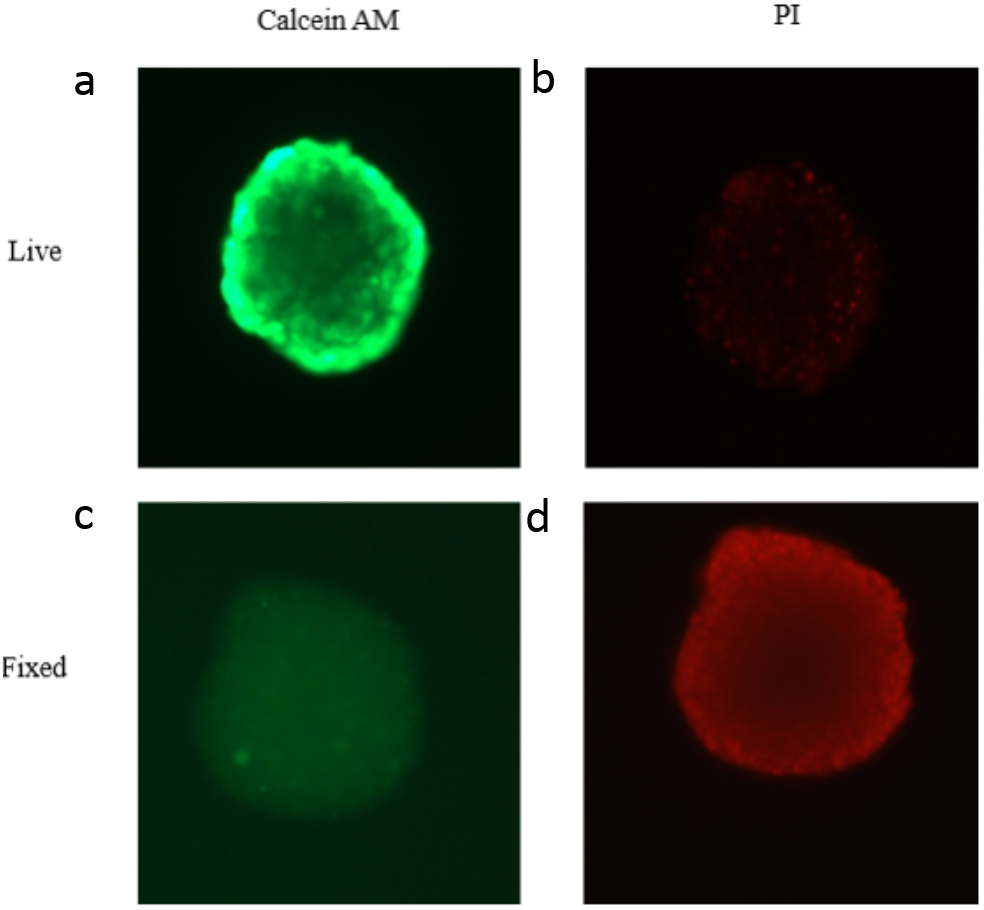
Viability assessment of the HepaRG cells in the spheroid. HepaRG spheroid is dyed with **(A)** Calcein-AM and **(B)** PI dyes for the evaluation of viability of cells. Then, spheroid is fixed with 70% ethanol for 10min and dyed with **(C)** Calcein-AM and **(C)** PI.

### 3D Spheroid Culture

Cell culture is one of the most important techniques for molecular and genetic analysis in the field. Most common culturing method of the cells is 2D monolayer culture which forces the cells to attach and grow on a flat surface. Cells in the body that grows in the 3D environment changes their metabolism and function when they are cultured as monolayers, since their interactions with other cells and extracellular matrix gets limited. Also, their gene expression profiles changes on 2D culture which as a result changes the response that the cells will give upon exposure to a toxic agent. 3D cell culture is the best way of mimicking the cell microenvironment. In 3D culture, the cells will give response against toxic agents more similar to in vivo. Therefore, we decided to generate 3D spheroids from HepaRG cells for the collecting mRNA for the RNA-seq analysis. We used agarose-based spheroid culture method for the generation of spheroids. This technique allows the cells to attach each other instead of attaching the plate surface. After 4 days of incubation on agarose plates, spheroids form the spheroid shape. Assessment of the viability of the cells inside the spheroid is done with staining the spheroids with Calcein-AM which dyes the live cells. Then, the dead cells are dyed with propidium iodide (PI) dye. Both staining showed that the cells inside the spheroid is mostly alive (**Figure 1**).

### Differential Expression

RNA-seq raw data was analysed by using Hisat, StringTie and Ballgown.^33^ Transcriptomic analysis provided detailed information on how HepaRG cells respond to nanoparticle cytotoxicity and which genes significantly changed their expression. Venn diagrams were generated to display the number of common differentially expressed genes. (**Figure 4B**) According to the results, 706, 839, 830 and 657 genes were differentially expressed with a p<0.05 in cells that were treated with AuNP, AgNP, TiO2NP and QDs, respectively. Among them 430 upregulated genes were identified for AuNP, 578 were identified for AgNP, 481 were identified for TiO2NP and 359 were identified for QDs; where 301 genes for AuNP, 283 genes for AgNP, 378 genes for TiO2NP and 319 genes for QD were found to be downregulated. (**Figure 4A**) Volcano plots shows the changes in the expression profiles of the cells exposed to nanoparticles of each material. (**Figure 4C-F**) The results revealed that nanoparticle exposure caused dramatic changes in the global gene transcription profile. Among all 4 types of nanoparticles, quantum dot was the most toxic type that generated dramatic changes in the gene expression profiles. Similarly, quantum dot had the smallest group of common DEGs with the other 3 groups. Gold, silver and titanium has 144 common genes upregulated, but the number of common genes for all 4 NP is 38, which is decreased by QD. This means that, the toxicity caused by quantum dots activated different pathways than the other 3 NPs, in other words its toxicity mechanism might be different than the other metal nanoparticles. Moreover, there were 293, 412, 302, 456 DEGs specific for only the nanoparticle type AuNP, AgNP, TiO_2_NP and QD, respectively.

**Figure 4.**
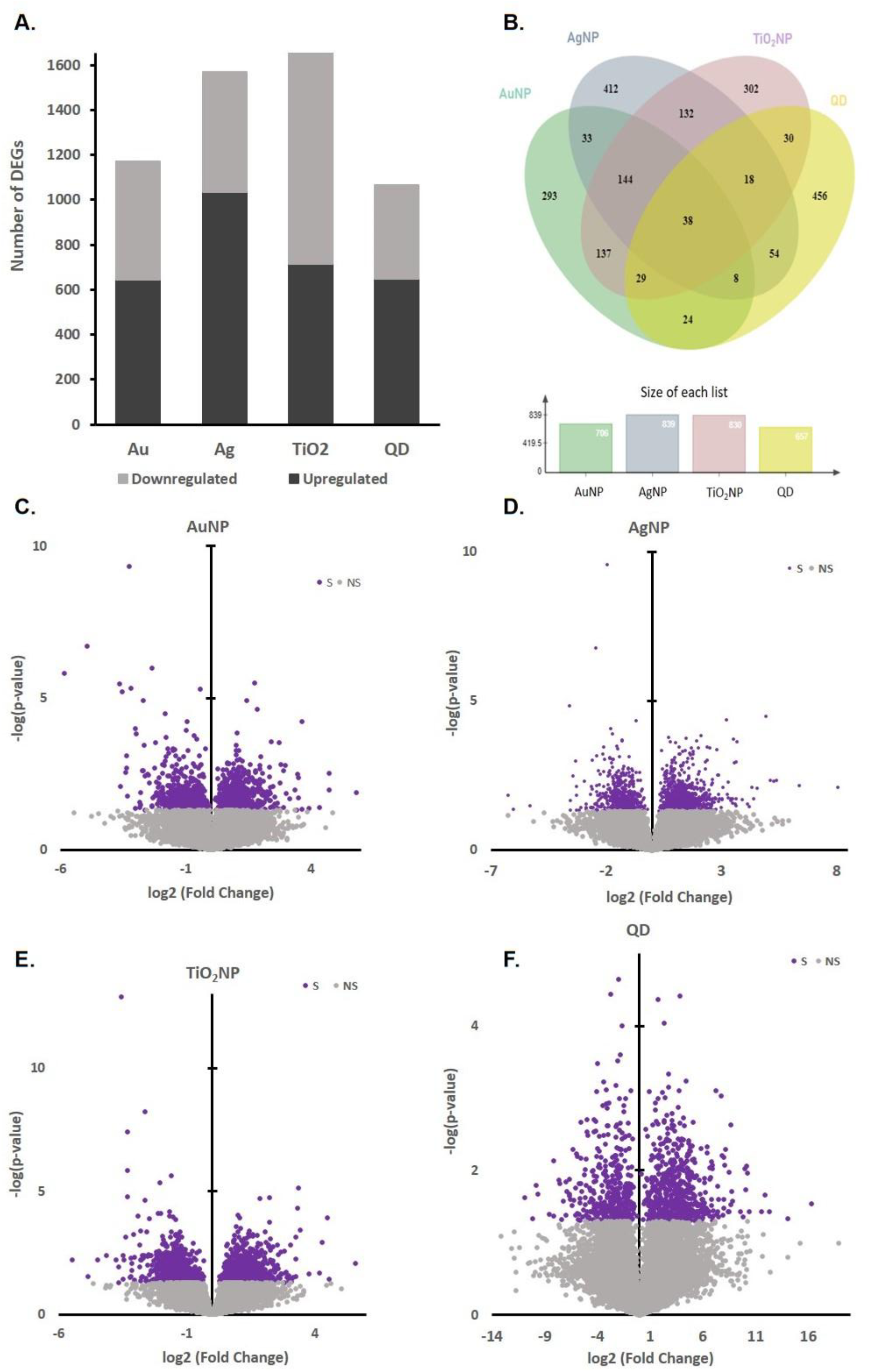
Changes in gene expression profiles of nanoparticle exposed cells **A)** Bar graph shows the number up upregulated (dark grey) and downregulated (light grey) genes in each transcriptome produced in response to each type of NPs. **B)** Venn diagram showing the number of DEGs in each transcriptome produced in response to each NP (P < 0.05). Volcano plot of gene expression profiles of the HepaRG cells **C)** gold vs. control **D)** silver vs. control **E)** titanium vs. control **F)** quantum dot vs. control represented with log2 (FC) and −log(p-value). Significant genes (purple) has a p-val<0.05, and non-significant genes has a p-val>0.05.

For the proceeding analysis, we set the criteria of FC>2 and p<0.05 for the determination of the DEGs. According to this criteria, among all 4 NPs, we found that 27 differentially expressed genes (DEGs) with FC>2 and p<0.05 were common. Among them 15 were upregulated and 6 were downregulated significantly. (**Supplementary Figure 1**) The rest were either upregulated or downregulated in different nanoparticles.

### Functional analysis of DEGs

In genome wide expression studies, many DEGs are found by data analysis and they need to be categorized to relate them to pathways or biological processes. In order to determine the function of DEGs and metabolic pathway enrichment, Gene Ontology (GO) enrichment analysis was performed on 27 DEGs (p<0.05 and FC>2) in each group. GO analysis was done to find the enrichment of the DEGs in three categories: biological process, cellular compartment and molecular function. In biological process category 96, 105, 156 and 115 terms were enriched in AuNP, AgNP, TiO2 and QD, respectively. **(Table 1)** However not all of them are common for all 4 types of NPs, where only 10 biological processes terms are significantly enriched for the common DEGs for all NPs.

**Table 1:** Number of GO terms and KEGG Pathways enriched in DEGs for each NP type.

| Nanoparticle | Biological Processes | Molecular Function | Cellular Compartment | Enriched KEGG Pathways |
| --- | --- | --- | --- | --- |
| Gold | 96 | 41 | 56 | 20 |
| Silver | 105 | 44 | 73 | 25 |
| Titanium | 156 | 97 | 72 | 29 |
| QD | 115 | 45 | 70 | 33 |
| Common DEGs for all 4 | 10 | 6 | 9 | 0 |

Gene ontology analysis showed that the enriched terms that are most relevant terms indicating nanoparticle toxicity were shown in the bar graph according to their p-values. (**Figure 5**) Many upregulated genes showed enrichment in similar pathways that are related to cellular response to metal ions, negative regulation of cell proliferation and apoptosis. Nanoparticle treatment induced expression of many genes that functions in the metal metabolism in the cell such as cellular iron ion homeostasis (GO:0006879), cellular response to zinc ion (GO:0071294), cellular copper ion homeostasis (GO:0006878) etc. Both AuNP and TiO2NP induced expression of BOLA2B, BOLA3, CP and ISCU which enriched the terms “iron ion homeostasis” (GO:0006879) and “iron-sulfur cluster assembly” (GO:0016226). AgNP induced the expression of metallothioneins MT1E, MT1F, MT1X and MT2A which enriched the terms “response to metal ion” (GO:0071248) and “detoxification of copper ion” (GO:0010273) where QD only induced the expression of MT2A. QDs also induced the expression of ATP2B1, ANO6, BEST1, P2RX4 and TTYH3 genes which function in the ion channels and enriched the term “ion transmembrane transport” (GO:0034220). In all of the nanoparticles the biological term “positive regulation of canonical NF-kappaB signal transduction” (GO:0043123) was enriched.

**Figure 5.**
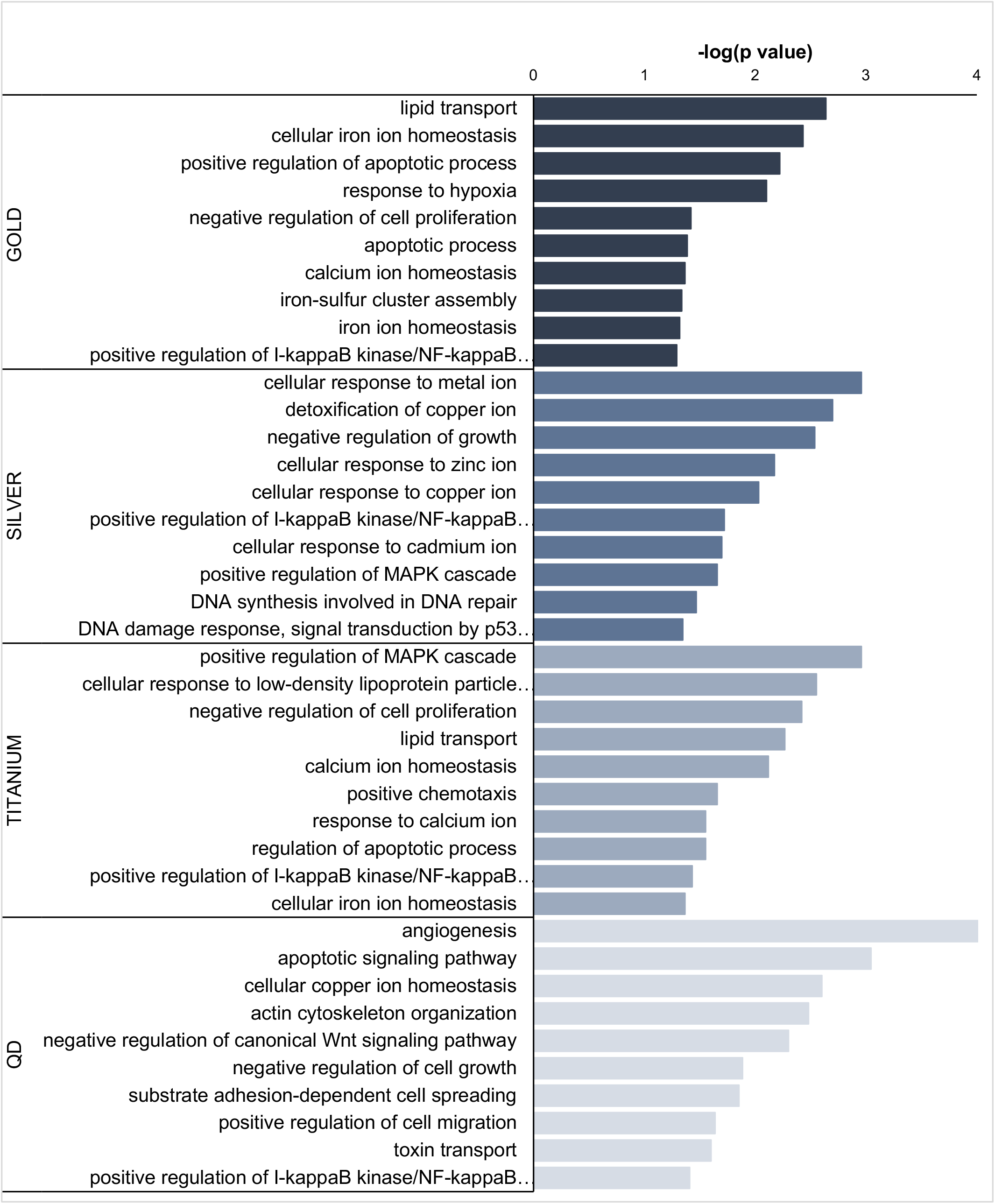
GO enrichment analysis of DEGs (p-val<0.05 and FC>2) for nanoparticles of each material. The ontology terms are listed in the bar graph according to their p-value. The x axis represents the –log(p-value) for each term, and the y axis represents the GO terms that were enriched for the common DEGs.

Gene ontology analysis of the common 27 DEGs showed that the top biological process enriched was cholesterol metabolic process (GO:0008203), top enriched molecular function GO term was chloride channel activity (GO:0005254), and the Golgi apparatus (GO:0005794) was the most enriched cellular component term. (**Supplementary Figure 2**). The other enriched biological processes clustered mainly in negative regulation of cell proliferation and chemotaxis of immune cells. GO analysis of the common DEGs showed that the NPs have disrupted the cellular cholesterol homeostasis of the cells where APP, APOL1 and CYP27A1 genes enriched the biological process terms “cholesterol metabolic process” (GO:0008203) and “lipoprotein metabolic process” (GO:0042157). (Supplementary Table 2) Shown that cation chloride metabolism was activated upon oxidative stress^34^, the terms “chloride transmembrane transport” (GO:1902476) and “chloride transport” (GO:0006821) was enriched upon nanoparticle caused toxicity in the cells.

KEGG pathway analysis showed that the upregulated pathways were related to oxidative stress, carcinogenesis, infection, and other diseases. One of the pathways that was upregulated in DEGs is the pathway that is related with Parkinson’s disease. **(Figure 6)** It was previously shown that nanoparticle might induce necrosis in cells causing Parkinson like neurobehavior in cells.^35^ This supports the results that are related to Parkinson’s disease and neurodegeneration.

**Figure 6.**
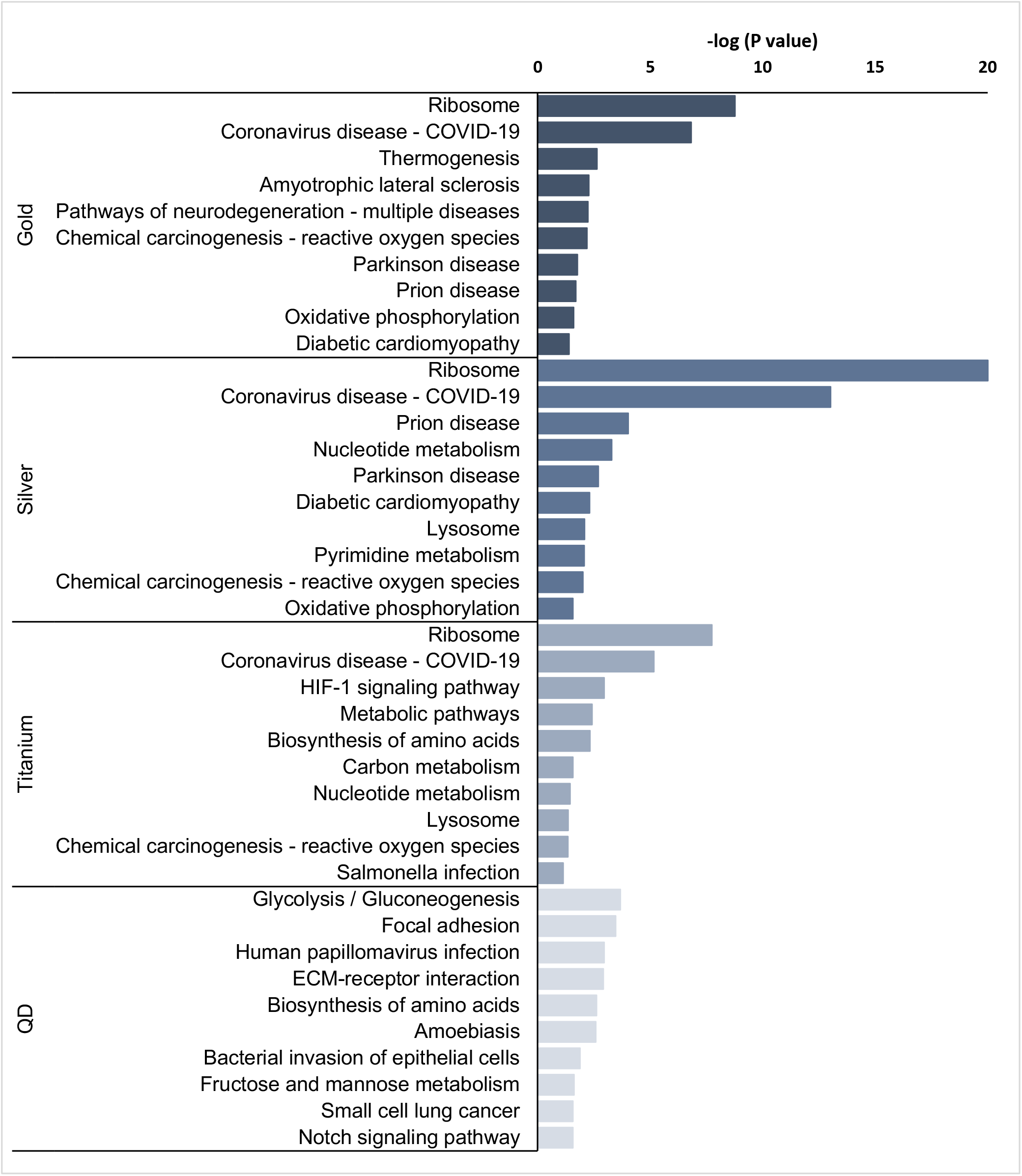
Top enriched KEGG pathways for the differentially upregulated genes in NP treated cells (P<0.05 and FC>2).

### Upregulated Genes

Upregulated genes common for all 4 nanoparticles were investigated. The genes that are related to toxicity response of the cells were chosen and the fold changes was shown in all of the nanoparticles. (**Table 2**) All nanoparticles induced overexpression of NDRG1 gene which is a tumor suppressor involved in stress response. ^36^ (**Table 2**) It is a membrane binding protein which binds to metal ions. CYP27A1 functions in cholesterol metabolism which has a tumor-suppressing effect. ^37^ Oncogenes were also upregulated such as IGFBP6^38,39^, PKM^40^ and TSPAN4^41^ which promote tumor cell proliferation and survival.

**Table 2:**
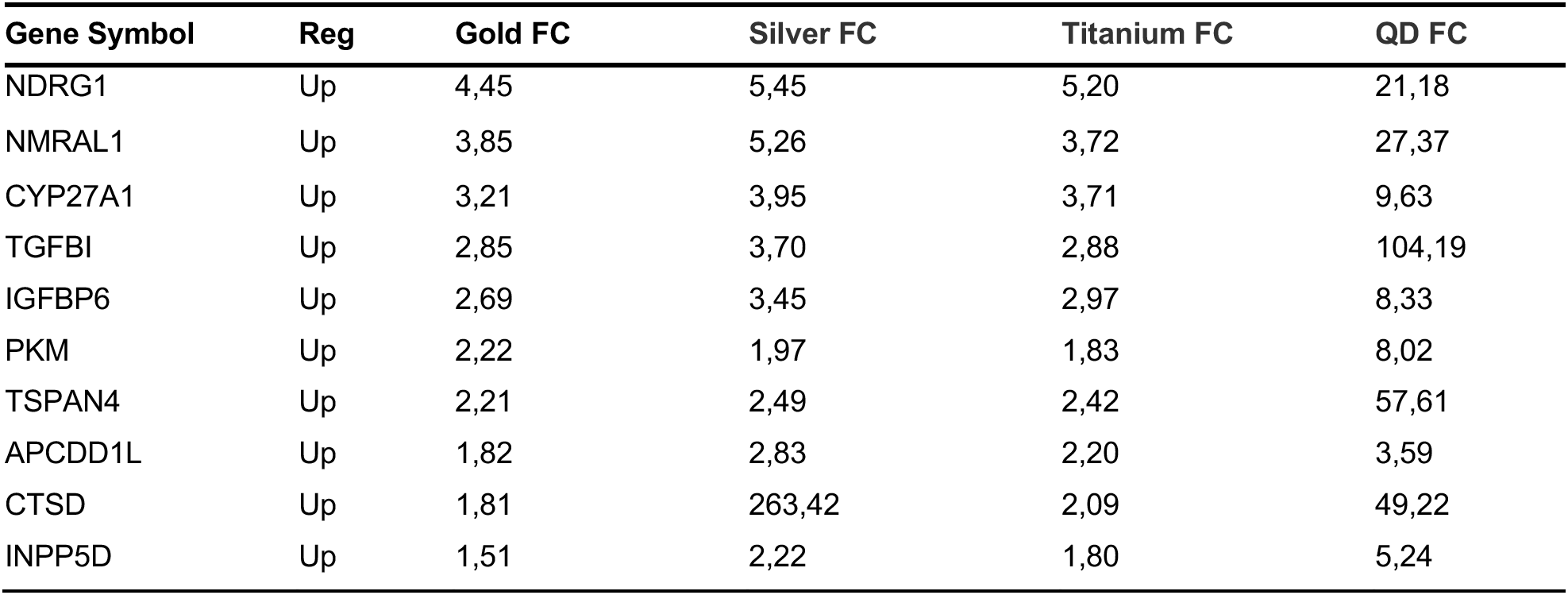
Common upregulated genes that are related to toxicity response of the cells.

## DISCUSSION

Nanoparticles were shown to cause cytotoxicity on cells.^42^ The small size of the nanoparticles enables them to penetrate into the cells and interact with molecules in the cells causing oxidative stress, DNA damage and inflammation.^43,44^ Our transcriptomic analysis demonstrated that the concentrations that causes viability loss=20% in AuNP, AgNP, TiO_2_NP and QDs caused cytotoxicity to HepaRG cells. As this study shows that all nanoparticles of different materials caused cytotoxicity on cells, it also demonstrated that the mechanism of cytotoxicity was different for each material. Number of common differentially expressed genes was higher for AuNP, AgNP and TiO_2_NP. They shared more DEGs between them as compared to QDs which decreased the number of common DEGs that we took for investigation. Also, the volcano plots showed that the most dramatic change in the expression profiles was seen in the cells that were treated with QDs. This might tell us that the mechanism of cytotoxicity of QDs is different than the other 3 nanoparticles.

Treatment with metal nanoparticles, the cells are exposed to metal ions since theories suggest that the nanoparticles release metal ions that disrupt cation homeostasis in cells, leading to cellular damage.^45,46^ So, the activation of pathways related to cation homeostasis is not surprising and shows that the cells are responding to these metal ions. Our results showed that the treatment with AuNP resulted in the increased expression of “cellular iron ion homeostasis” and “calcium ion homeostasis”. Treatment with AgNP resulted in the increased expression “detoxification of copper ion” and metal ion response pathways such as cellular response to zinc, copper and cadmium ion. TiO_2_NP caused the activation “calcium ion homeostasis” and “cellular response to iron ion” pathways. QDs activated the pathways “cellular copper ion homeostasis” and “toxin transport”. These results indicate that the cells are exposed with the metal ions when treated with nanoparticles, and these metal ions are activating the pathways that function for the cellular ion homeostasis.

Nanoparticles activated the expression of many genes that promote apoptosis. Enrichment of the apoptosis related pathways indicates the levels of the cytotoxicity that caused by the nanoparticles. Among common DEGs NDRG1, OPN3, PKM, INPP5D are the genes that are related to apoptosis. It is shown that the overexpression of NDRG1 in cancer cells decreased the survival and induces apoptosis. ^47^ APP and PIGT genes together enriched the term that “neuron apoptotic process” which are upregulated in all nanoparticle caused cytotoxicity. APP was shown to induce apoptosis in neurons.^48^

Nanoparticles induce increase in the intracellular ROS production because of their strong oxidation ability.^49^ Excess ROS production results in damage in DNA, organelles, cell membrane and cause lipid peroxidation and apoptosis.^50^ When ROS levels rises, cells activate the detoxification pathways.^51^ One of these pathways is MAPK signaling^52^ which is seen in the AgNP and TiO_2_NP treated cells. MAPK cascade is one of the most important signaling pathways that functions as a transmitters of the stress causing stimuli especially ROS.^53^ IGFBP6, upregulated by the cytotoxicity of all NPs, is found to activate stress activated MAPK cascade.

NF-kappaB pathway regulates the immune and inflammatory responses.^54,55^ It induces the expression of many cytokines and chemokines. Also it functions in the promotion of cell survival by inhibiting apoptosis related pathways. This function of NF-kappaB pathway is used by cancer cells to avoid cell death against drugs, radiation and cytokines.^56^ The cells in our study activated the NF-kappaB pathway upon treatment with all of the nanoparticles independent of their material. The genes that enriched this term are not common however enrichment of this term in all of the nanoparticle types may indicate that the cells were under stress and they were trying to survive.

Carcinogenic potential of the nanoparticles was reported previously in many studies because of their genotoxic effects on cells.^57,58,59^ The mechanism that NPs might induce the cells to evolve to tumor cells is the ability to cause DNA damage. In this study, we found that the AgNP activates pathways related to “DNA synthesis involved in DNA repair” and “DNA damage response, signal transduction by p53 class mediator resulting in cell cycle arrest” were activated. Also, the expression of the genes related to cell migration such as TGFBI and IGFBP6 were upregulated. TGFBI and IGFBP6 were shown to promote the migration and invasion of cancer cells.^60,61^ An enzyme that functions in glycolysis, PKM was upregulated in all of the nanoparticle treated cells. With the knowledge that the cancer cells rely on glycolysis, PKM induces the growth and migration of the cancer cells, where knockdown of PKM inhibited the growth and migration of the breast cancer cells. ^62^ NP toxicity also induced the expression of OPN3 which is a cell surface photoreceptor, induced tumorigenesis, EMT and metastasis of cancer cells.^63,65^ It was also shown to increase the risk of recurrence in cancer patients.

## Data Availability

All of the data is available upon request including genomic data (raw or processed), gene annotation are given as tables in the supplementary data.

## Supplementary Data Statement

Supplementary Data are available at NAR online.

## MATERIAL AND METHODS

### Reagents

HepaRG cells (Cat.No. HPRGC10), Pen/Strep and FBS were purchased from Thermo Scientific. William’s E Medium and hydrocortisone were purchased from Sigma Aldrich. Human insulin was purchased from pharmacy (Humulin N, 100U/mL).

### Cell Culture

HepaRG cells were purchased from Thermo Scientific (Cat.No. HPRGC10). Cells were cultured in William’s E Medium supplemented with 10% FBS, 100 U/ml Pen/Strep, 2mM L-glutamine, 32 mU/mL human insulin and 20 µg/mL hydrocortisone. Cells were cultured in T25 flask until 80% confluency changing medium every 2 days.

### Nanoparticle Characterization

Nanoparticles obtained were buffer exchanged into full William’s E Medium and then characterized by adding 100µL of each nanoparticle to 96 well plates and taking the absorbance measurements within 350nm-750nm wavelength spectrum in a microplate reader. TEM images were obtained by adding 10µL of nanoparticles to a grid, then washing the grid with 10µL of water for 3 times.

### MTT Assay

HepaRG cells that were cultured in T25 flask were collected with trypsin, then seeded into the 96 well plated in a density of 30,000 cells/well and cultured for 1 day. The other day, increasing concentrations of nanoparticles were added on to the cells in fresh medium. Cells and nanoparticles were incubated together for 24hr and the medium was changed with 10% MTT containing medium. After 4h of incubation, medium was discarded and the purple crystals were dissolved in 100µL DMSO. Measurements were taken with microplate reader at 560nm.

### Agarose method for spheroid formation

96 well plates were coated with 50µL sterile 1.5% cell culture grade agarose dissolved in PBS. Then HepaRG cells were seeded into the plate in a density of 3,000 cells/well in 100µL medium. Cells were cultured for 3 days for the spheroids to form. At day 3, the spheroids were collected and put into a 6 well plate coated with 2mL 1.5% agarose. Then the appropriate concentrations of nanoparticles were added on to the spheroids, and incubated for 24 hr. Next day, the spheroids were collected by centrifugation and were subjected to RNA isolation (M&N, NucleoSpin RNA, Cat.No.740955.50) procedure according to the manufacturer’s protocol.

### High-throughput transcriptomic sequencing

Total RNAs that are isolated from HepaRG spheroids (treated with gold nanoparticles (3nm, 15nm, 80nm), silver nanoparticles (12nm, 40nm, 80nm), titanium nanoparticles (25nm, 100nm, 300nm) and 2 quantum dots (CdZnSe/CdZnS/ZnS and InP/ZnS) for 24 hours, (each nanoparticle and control group has 2 replicates). Total RNA was extracted using NucleoSpin (Macherey-Nagel™ NucleoSpin™ RNA, Mini Kit) following the manufacturer’s protocol. The quantity and quality of RNA were examined by Termo ScientifcTM NanoDropTM 8000 Spectrophotometer. Library sequencing was performed on an Illumina Hiseq. 4000 platform, to create paired-end reads with a length of 150 bp.

### Bioinformatics analyses

The quality control of RNA-Seq data was conducted using the FastQC with default parameters. The RNA-seq data was analyzed according to the protocol that was proposed by Pertea et al. ^33^ Clean paired-end reads were aligned to the human reference genome sequence, GRCz10 version31, using Hisat2. To identify differential expression genes (DEGs) between the control and treated groups the expression levels are calculated by StringTie. Ballgown was used for the differential expression analysis. The DEGs between two groups were selected based on the p-value<0.05 and fold change>2. To understand the functions of the differentially expressed genes, gene ontology (GO) functional enrichment and KEGG pathway analysis was done with NIH DAVID Bioinformatics tool (https://david.ncifcrf.gov/tools.jsp).

### Statistical Analysis

The Ballgown calculated p-value was used to determine the significant DEGs. A p-val <0.05 and fold change >2 were used as the threshold to determine the statistical significantly upregulated genes. For GO enrichment analysis, the Benjamini-Hochberg p-value and the P-value (<0.05) was used to determine the significant enrichment of the gene sets. For KEGG enrichment analysis, a p-val <0.05 was used as the threshold to judge the significant enrichment of the gene sets. A P-value<0.05 was considered statistically significant. The DEGs between two groups were selected based on the following criteria: 1) the fold change was greater than 2, and 2) the p-value was less than 0.05.

## References

(1) Bystrzejewska-Piotrowska, G.; Golimowski, J.; Urban, P. L. Nanoparticles: Their Potential Toxicity, Waste and Environmental Management. Waste Manag. 2009, 29 (9), 2587–2595. 10.1016/j.wasman.2009.04.001.

(2) Khan, Y.; Sadia, H.; Ali Shah, S. Z.; Khan, M. N.; Shah, A. A.; Ullah, N.; Ullah, M. F.; Bibi, H.; Bafakeeh, O. T.; Khedher, N. B.; Eldin, S. M.; Fadhl, B. M.; Khan, M. I. Classification, Synthetic, and Characterization Approaches to Nanoparticles, and Their Applications in Various Fields of Nanotechnology: A Review. Catalysts 2022, 12 (11), 1386. 10.3390/catal12111386.

(3) Ealia, S. A. M.; Saravanakumar, M. P. A Review on the Classification, Characterisation, Synthesis of Nanoparticles and Their Application. IOP Conf. Ser. Mater. Sci. Eng. 2017, 263 (3), 032019. 10.1088/1757-899X/263/3/032019.

(4) Patil, S. S.; Nitave, D. S. A. A REVIEW ON THE NANOPARTICLES. World J. Pharm. Res.

(5) Khan, I.; Saeed, K.; Khan, I. Nanoparticles: Properties, Applications and Toxicities. Arab. J. Chem. 2019, 12 (7), 908–931. 10.1016/j.arabjc.2017.05.011.

(6) López-Serrano, A.; Olivas, R. M.; Landaluze, J. S.; Cámara, C. Nanoparticles: A Global Vision. Characterization, Separation, and Quantification Methods. Potential Environmental and Health Impact. Anal. Methods 2013, 6 (1), 38–56. 10.1039/C3AY40517F.

(7) De, M.; Ghosh, P. S.; Rotello, V. M. Applications of Nanoparticles in Biology. Adv. Mater. 2008, 20 (22), 4225–4241. 10.1002/adma.200703183.

(8) Khoshnevisan, K.; Daneshpour, M.; Barkhi, M.; Gholami, M.; Samadian, H.; Maleki, H. The Promising Potentials of Capped Gold Nanoparticles for Drug Delivery Systems. J. Drug Target. 2018, 26 (7), 525–532. 10.1080/1061186X.2017.1387790.

(9) Sardar, R.; Funston, A. M.; Mulvaney, P.; Murray, R. W. Gold Nanoparticles: Past, Present, and Future. Langmuir 2009, 25 (24), 13840–13851. 10.1021/la9019475.

(10) Su, S.; Zuo, X.; Pan, D.; Pei, H.; Wang, L.; Fan, C.; Huang, W. Design and Applications of Gold Nanoparticle Conjugates by Exploiting Biomolecule–Gold Nanoparticle Interactions. Nanoscale 2013, 5 (7), 2589. 10.1039/c3nr33870c.

(11) Kedziora, A.; Gorzelańczyk, K.; Bugla-Płoskońska, G. Positive and Negative Aspects of Silver Nanoparticles Usage. Biol Int 2013, 53, 67–76.

(12) Ravindran, A.; Chandran, P.; Khan, S. S. Biofunctionalized Silver Nanoparticles: Advances and Prospects. Colloids Surf. B Biointerfaces 2013, 105, 342–352. 10.1016/j.colsurfb.2012.07.036.

(13) Çeşmeli, S.; Biray Avci, C. Application of Titanium Dioxide (TiO2) Nanoparticles in Cancer Therapies. J. Drug Target. 2019, 27 (7), 762–766. 10.1080/1061186X.2018.1527338.

(14) Drbohlavova, J.; Adam, V.; Kizek, R.; Hubalek, J. Quantum Dots — Characterization, Preparation and Usage in Biological Systems. Int. J. Mol. Sci. 2009, 10 (2), 656–673. 10.3390/ijms10020656.

(15) McNamara, K.; Tofail, S. A. M. Nanoparticles in Biomedical Applications. Adv. Phys. X 2017, 2 (1), 54–88. 10.1080/23746149.2016.1254570.

(16) Arruebo, M.; Fernández-Pacheco, R.; Ibarra, M. R.; Santamaría, J. Magnetic Nanoparticles for Drug Delivery. Nano Today 2007, 2 (3), 22–32. 10.1016/S1748-0132(07)70084-1.

(17) Saltepe, B.; Bozkurt, E. U.; Hacıosmanoğlu, N.; Şeker, U. Ö. Ş. Genetic Circuits To Detect Nanomaterial Triggered Toxicity through Engineered Heat Shock Response Mechanism. ACS Synth. Biol. 2019, 8 (10), 2404–2417. 10.1021/acssynbio.9b00291.

(18) Seaton, A.; Tran, L.; Aitken, R.; Donaldson, K. Nanoparticles, Human Health Hazard and Regulation. J. R. Soc. Interface 2010, 7 Suppl 1 (Suppl 1), S119–129. 10.1098/rsif.2009.0252.focus.

(19) Brown, L. M.; Collings, N.; Harrison, R. M.; Maynard, A. D.; Maynard, R. L.; Donaldson, K.; Stone, V.; Gilmour, P. S.; Brown, D. M.; MacNee, W. Ultrafine Particles: Mechanisms of Lung Injury. Philos. Trans. R. Soc. Lond. Ser. Math. Phys. Eng. Sci. 2000, 358 (1775), 2741–2749. 10.1098/rsta.2000.0681.

(20) Love, S. A.; Maurer-Jones, M. A.; Thompson, J. W.; Lin, Y.-S.; Haynes, C. L. Assessing Nanoparticle Toxicity. Annu. Rev. Anal. Chem. Palo Alto Calif 2012, 5, 181–205. 10.1146/annurev-anchem-062011-143134.

(21) Smolkova, B.; El Yamani, N.; Collins, A. R.; Gutleb, A. C.; Dusinska, M. Nanoparticles in Food. Epigenetic Changes Induced by Nanomaterials and Possible Impact on Health. Food Chem. Toxicol. 2015, 77, 64–73. 10.1016/j.fct.2014.12.015.

(22) Commissioner, O. of the. Considering Whether an FDA-Regulated Product Involves the Application of Nanotechnology. U.S. Food and Drug Administration, 2019. https://www.fda.gov/regulatory-information/search-fda-guidance-documents/considering-whether-fda-regulated-product-involves-application-nanotechnology (accessed 2023-05-26).

(23) Martínez, G.; Merinero, M.; Pérez-Aranda, M.; Pérez-Soriano, E. M.; Ortiz, T.; Begines, B.; Alcudia, A. Environmental Impact of Nanoparticles’ Application as an Emerging Technology: A Review. Materials 2020, 14 (1), 166. 10.3390/ma14010166.

(24) Anthérieu, S.; Chesné, C.; Li, R.; Guguen-Guillouzo, C.; Guillouzo, A. Optimization of the HepaRG Cell Model for Drug Metabolism and Toxicity Studies. Toxicol. In Vitro 2012, 26 (8), 1278–1285. 10.1016/j.tiv.2012.05.008.

(25) Marion, M.-J.; Hantz, O.; Durantel, D. The HepaRG Cell Line: Biological Properties and Relevance as a Tool for Cell Biology, Drug Metabolism, and Virology Studies. In Hepatocytes: Methods and Protocols; Maurel, P., Ed.; Methods in Molecular Biology; Humana Press: Totowa, NJ, 2010; pp 261–272. 10.1007/978-1-60761-688-7_13.

(26) Gunti, S.; Hoke, A. T. K.; Vu, K. P.; London, N. R. Organoid and Spheroid Tumor Models: Techniques and Applications. Cancers 2021, 13 (4), 874. 10.3390/cancers13040874.

(27) Tseng, H.; Gage, J. A.; Shen, T.; Haisler, W. L.; Neeley, S. K.; Shiao, S.; Chen, J.; Desai, P. K.; Liao, A.; Hebel, C.; Raphael, R. M.; Becker, J. L.; Souza, G. R. A Spheroid Toxicity Assay Using Magnetic 3D Bioprinting and Real-Time Mobile Device-Based Imaging. Sci. Rep. 2015, 5 (1), 13987. 10.1038/srep13987.

(28) Flampouri, E.; Imar, S.; OConnell, K.; Singh, B. Spheroid-3D and Monolayer-2D Intestinal Electrochemical Biosensor for Toxicity/Viability Testing: Applications in Drug Screening, Food Safety, and Environmental Pollutant Analysis. ACS Sens. 2019, 4 (3), 660–669. 10.1021/acssensors.8b01490.

(29) Fey, S. J.; Wrzesinski, K. Determination of Drug Toxicity Using 3D Spheroids Constructed From an Immortal Human Hepatocyte Cell Line. Toxicol. Sci. 2012, 127 (2), 403–411. 10.1093/toxsci/kfs122.

(30) Han, Y.; Gao, S.; Muegge, K.; Zhang, W.; Zhou, B. Advanced Applications of RNA Sequencing and Challenges. Bioinforma. Biol. Insights 2015, 9s1, BBI.S28991. 10.4137/BBI.S28991.

(31) Kukurba, K. R.; Montgomery, S. B. RNA Sequencing and Analysis. Cold Spring Harb. Protoc. 2015, 2015 (11), pdb.top084970. 10.1101/pdb.top084970.

(32) Pertea, M.; Kim, D.; Pertea, G. M.; Leek, J. T.; Salzberg, S. L. Transcript-Level Expression Analysis of RNA-Seq Experiments with HISAT, StringTie and Ballgown. Nat. Protoc. 2016, 11 (9), 1650–1667. 10.1038/nprot.2016.095.

(33) Piechotta, K.; Lu, J.; Delpire, E. Cation Chloride Cotransporters Interact with the Stress-Related Kinases Ste20-Related Proline-Alanine-Rich Kinase (SPAK) and Oxidative Stress Response 1 (OSR1)*. J. Biol. Chem. 2002, 277 (52), 50812–50819. 10.1074/jbc.M208108200.

(34) Sadeghi, L.; Babadi, V. Y.; Tanwir, F. Manganese Dioxide Nanoparticle Induces Parkinson like Neurobehavioral Abnormalities in Rats. Bratisl. Med. J. 2018, 119 (06), 379–384. 10.4149/BLL_2018_070.

(35) Fang, B. A.; Kovačević, Ž.; Park, K. C.; Kalinowski, D. S.; Jansson, P. J.; Lane, D. J. R.; Sahni, S.; Richardson, D. R. Molecular Functions of the Iron-Regulated Metastasis Suppressor, NDRG1, and Its Potential as a Molecular Target for Cancer Therapy. Biochim. Biophys. Acta BBA - Rev. Cancer 2014, 1845 (1), 1–19. 10.1016/j.bbcan.2013.11.002.

(36) Liang, Z.; Jiao, W.; Wang, L.; Chen, Y.; Li, D.; Zhang, Z.; Zhang, Z.; Liang, Y.; Niu, H. CYP27A1 Inhibits Proliferation and Migration of Clear Cell Renal Cell Carcinoma via Activation of LXRs/ABCA1. Exp. Cell Res. 2022, 419 (1), 113279. 10.1016/j.yexcr.2022.113279.

(37) Oliva, C. R.; Halloran, B.; Hjelmeland, A. B.; Vazquez, A.; Bailey, S. M.; Sarkaria, J. N.; Griguer, C. E. IGFBP6 Controls the Expansion of Chemoresistant Glioblastoma through Paracrine IGF2/IGF-1R Signaling. Cell Commun. Signal. 2018, 16 (1), 61. 10.1186/s12964-018-0273-7.

(38) Zhao, C.; Zhu, X.; Wang, G.; Wang, W.; Ju, S.; Wang, X. Decreased Expression of IGFBP6 Correlates with Poor Survival in Colorectal Cancer Patients. Pathol. - Res. Pract. 2020, 216 (5), 152909. 10.1016/j.prp.2020.152909.

(39) Zahra, K.; Dey, T.; Ashish; Mishra, S. P.; Pandey, U. Pyruvate Kinase M2 and Cancer: The Role of PKM2 in Promoting Tumorigenesis. Front. Oncol. 2020, 10.

(40) Zheng, Y.; Lang, Y.; Qi, B.; Li, T. TSPAN4 and Migrasomes in Atherosclerosis Regression Correlated to Myocardial Infarction and Pan-Cancer Progression. Cell Adhes. Migr. 2023, 17 (1), 14–19. 10.1080/19336918.2022.2155337.

(41) Xiong, P.; Huang, X.; Ye, N.; Lu, Q.; Zhang, G.; Peng, S.; Wang, H.; Liu, Y. Cytotoxicity of Metal-Based Nanoparticles: From Mechanisms and Methods of Evaluation to Pathological Manifestations. Adv. Sci. 2022, 9 (16), 2106049. 10.1002/advs.202106049.

(42) Xia, T.; Kovochich, M.; Liong, M.; Mädler, L.; Gilbert, B.; Shi, H.; Yeh, J. I.; Zink, J. I.; Nel, A. E. Comparison of the Mechanism of Toxicity of Zinc Oxide and Cerium Oxide Nanoparticles Based on Dissolution and Oxidative Stress Properties. ACS Nano 2008, 2 (10), 2121–2134. 10.1021/nn800511k.

(43) Chen, P. Detection of DNA Damage Response Caused by Different Forms of Titanium Dioxide Nanoparticles Using Sensor Cells. J. Biosens. Bioelectron. 2012, 03 (05). 10.4172/2155-6210.1000129.

(44) Ma, H.; Guo, L.; Zhang, H.; Wang, Y.; Miao, Y.; Liu, X.; Peng, M.; Deng, X.; Peng, Y.; Fan, H. The Metal Ion Release of Manganese Ferrite Nanoparticles: Kinetics, Effects on Magnetic Resonance Relaxivities, and Toxicity. ACS Appl. Bio Mater. 2022, 5 (6), 3067–3074. 10.1021/acsabm.2c00338.

(45) Multi-omics approaches confirm metal ions mediate the main toxicological pathways of metal-bearing nanoparticles in lung epithelial A549 cells - Environmental Science: Nano (RSC Publishing). https://pubs.rsc.org/en/content/articlelanding/2018/en/c8en00071a (accessed 2023-08-31).

(46) Ma, W.; Na, M.; Tang, C.; Wang, H.; Lin, Z. Overexpression of N-myc Downstream-regulated Gene 1 Inhibits Human Glioma Proliferation and Invasion via Phosphoinositide 3-kinase/AKT Pathways. Mol. Med. Rep. 2015, 12 (1), 1050–1058. 10.3892/mmr.2015.3492.

(47) Chen, Y.; Liu, W.; McPhie, D. L.; Hassinger, L.; Neve, R. L. APP-BP1 Mediates APP-Induced Apoptosis and DNA Synthesis and Is Increased in Alzheimer’s Disease Brain. J. Cell Biol. 2003, 163 (1), 27–33. 10.1083/jcb.200304003.

(48) Manke, A.; Wang, L.; Rojanasakul, Y. Mechanisms of Nanoparticle-Induced Oxidative Stress and Toxicity. BioMed Res. Int. 2013, 2013, 942916. 10.1155/2013/942916.

(49) Yu, Z.; Li, Q.; Wang, J.; Yu, Y.; Wang, Y.; Zhou, Q.; Li, P. Reactive Oxygen Species-Related Nanoparticle Toxicity in the Biomedical Field. Nanoscale Res. Lett. 2020, 15 (1), 115. 10.1186/s11671-020-03344-7.

(50) He, L.; He, T.; Farrar, S.; Ji, L.; Liu, T.; Ma, X. Antioxidants Maintain Cellular Redox Homeostasis by Elimination of Reactive Oxygen Species. Cell. Physiol. Biochem. 2017, 44 (2), 532–553. 10.1159/000485089.

(51) Nahirnyj, A.; Livne-Bar, I.; Guo, X.; Sivak, J. M. ROS Detoxification and Proinflammatory Cytokines Are Linked by P38 MAPK Signaling in a Model of Mature Astrocyte Activation. PLOS ONE 2013, 8 (12), e83049. 10.1371/journal.pone.0083049.

(52) Jalmi, S.; Sinha, A. ROS Mediated MAPK Signaling in Abiotic and Biotic Stress-Striking Similarities and Differences. Front. Plant Sci. 2015, 6.

(53) Liu, T.; Zhang, L.; Joo, D.; Sun, S.-C. NF-κB Signaling in Inflammation. Signal Transduct. Target. Ther. 2017, 2, 17023. 10.1038/sigtrans.2017.23.

(54) Li, Q.; Verma, I. M. NF-κB Regulation in the Immune System. Nat. Rev. Immunol. 2002, 2 (10), 725–734. 10.1038/nri910.

(55) Luo, J.-L.; Kamata, H.; Karin, M. IKK/NF-κB Signaling: Balancing Life and Death – a New Approach to Cancer Therapy. J. Clin. Invest. 2005, 115 (10), 2625–2632. 10.1172/JCI26322.

(56) Ahamed, M.; Karns, M.; Goodson, M.; Rowe, J.; Hussain, S. M.; Schlager, J. J.; Hong, Y. DNA Damage Response to Different Surface Chemistry of Silver Nanoparticles in Mammalian Cells. Toxicol. Appl. Pharmacol. 2008, 233 (3), 404–410. 10.1016/j.taap.2008.09.015.

(57) Lim, H. K.; AshaRani, P. V.; Hande, M. P. Enhanced Genotoxicity of Silver Nanoparticles in DNA Repair Deficient Mammalian Cells. Front. Genet. 2012, 3.

(58) Klien, K.; Godnić-Cvar, J. Genotoxicity of Metal Nanoparticles: Focus on in Vivo Studies. Arh. Hig. Rada Toksikol. 2012, 63 (2), 133–145. 10.2478/10004-1254-63-2012-2213.

(59) Shang, D.; Liu, Y.; Yang, P.; Chen, Y.; Tian, Y. TGFBI-Promoted Adhesion, Migration and Invasion of Human Renal Cell Carcinoma Depends on Inactivation of von Hippel-Lindau Tumor Suppressor. Urology 2012, 79 (4), 966.e1–7. 10.1016/j.urology.2011.12.011.

(60) Fu, P.; Thompson, J. A.; Bach, L. A. Promotion of Cancer Cell Migration: AN INSULIN-LIKE GROWTH FACTOR (IGF)-INDEPENDENT ACTION OF IGF-BINDING PROTEIN-6*. J. Biol. Chem. 2007, 282 (31), 22298–22306. 10.1074/jbc.M703066200.

(61) Ma, C.; Zu, X.; Liu, K.; Bode, A. M.; Dong, Z.; Liu, Z.; Kim, D. J. Knockdown of Pyruvate Kinase M Inhibits Cell Growth and Migration by Reducing NF-kB Activity in Triple-Negative Breast Cancer Cells. Mol. Cells 2019, 42 (9), 628–636. 10.14348/molcells.2019.0038.

(62) Xu, C.; Wang, R.; Yang, Y.; Xu, T.; Li, Y.; Xu, J.; Jiang, Z. Expression of OPN3 in Lung Adenocarcinoma Promotes Epithelial-mesenchymal Transition and Tumor Metastasis. Thorac. Cancer 2020, 11 (2), 286–294. 10.1111/1759-7714.13254.

(63) Miyanaga, A.; Masuda, M.; Motoi, N.; Tsuta, K.; Nakamura, Y.; Nishijima, N.; Watanabe, S.; Asamura, H.; Tsuchida, A.; Seike, M.; Gemma, A.; Yamada, T. Whole-Exome and RNA Sequencing of Pulmonary Carcinoid Reveals Chromosomal Rearrangements Associated with Recurrence. Lung Cancer 2020, 145, 85–94. 10.1016/j.lungcan.2020.03.027.

