## Supplemental file for "Transcriptomic Implications of Toxic Effects of Nanoparticles on Metabolic Pathways of Liver Cells"

| **Gene ID** | **Symbol** | **Description** | **Function** |
| --- | --- | --- | --- |
| ENSG00000124155 | PIGT | Phosphatidylinositol Glycan Anchor Biosynthesis Class T | Component of the GPI transamidase complex.  Essential for transfer of GPI to proteins, particularly for formation of carbonyl intermediates |
| ENSG00000166165 | CKB | Creatine Kinase B | Reversibly catalyzes the transfer of phosphate between ATP and various phosphogens, creatine kinase isoenzymes play a central role in energy transduction in tissues with large, fluctuating energy demands, acts as a key regulator of adaptive thermogenesis as part of the futile creatine cycle |
| ENSG00000203875 | SNHG5 | Small Nucleolar RNA Host Gene 5 | RNA processing |
| ENSG00000181458 | TMEM45A | Transmembrane Protein 45A | Predicted to be integral component of membrane |
| ENSG00000104419 | NDRG1 | N-Myc Downstream Regulated 1 | Stress responses, hormone responses, cell growth, and differentiation, p53-mediated caspase activation and apoptosis, tumor suppressor, cell trafficking, p53/TP53-dependent mitotic spindle checkpoint |
| ENSG00000153406 | NMRAL1 | NmrA Like Redox Sensor 1 | Redox sensor protein, responses to changes in intracellular NADPH/NADP(+) levels, negatively regulates the activity of NF-kappaB in a ubiquitylation-dependent manner, cellular antiviral response |
| ENSG00000136295 | TTYH3 | Tweety Family Member 3 | chloride anion channels, Ca(2+) signal transduction |
| ENSG00000135929 | CYP27A1 | Cytochrome P450 Family 27 Subfamily A Member 1 | drug metabolism and synthesis of cholesterol, steroids and other lipids, cholesterol homeostasis, vitamin D biosynthesis |
| ENSG00000120708 | TGFBI | Transforming Growth Factor Beta Induced | cell adhesion, cell-collagen interactions, induced by transforming growth factor-beta and acts to inhibit cell adhesion |
| ENSG00000113163 | CERT1 | Ceramide Transporter 1 | ceramide intracellular transport |
| ENSG00000167779 | IGFBP6 | Insulin Like Growth Factor Binding Protein 6 | cell migration, positive regulation of stress-activated MAPK cascade, biomarker of breast cancer, in situ carcinoma, inhibit or stimulate the growth |
| ENSG00000152082 | MZT2B | Mitotic Spindle Organizing Protein 2B | protein binding |
| ENSG00000054277 | OPN3 | Opsin 3 | photoreception , regulates apoptosis via cytochrome c release, melanocyte survival |
| ENSG00000163710 | PCOLCE2 | Procollagen C-Endopeptidase Enhancer 2 | collagen binding activity; heparin binding activity; and peptidase activator activity, |
| ENSG00000067225 | PKM | Pyruvate Kinase M1/2 | Glycolysis (tumor cell proliferation and survival), regulation of transcription, cell proliferation, tumorigenesis, cell death of tumor cells |
| ENSG00000214063 | TSPAN4 | Tetraspanin 4 | signal transduction events that play a role in the regulation of cell development, activation, growth and motility |
| ENSG00000198768 | APCDD1L | APC Down-Regulated 1 Like | Wnt-protein binding activity, integral component of membrane |
| ENSG00000117984 | CTSD | Cathepsin D | pathogenesis of several diseases such as breast cancer and possibly Alzheimer disease |
| ENSG00000168918 | INPP5D | Inositol Polyphosphate-5-Phosphatase D | negatively regulating the PI3K pathways, negative regulator of B-cell antigen receptor signaling, cell-cell junctions, neutrophil migration, mediates activin/TGF-beta-induced apoptosis |

**Supplementary**

**Supplementary Table 1**: The common upregulated genes in nanoparticle exposed cells each compared with the

control group.

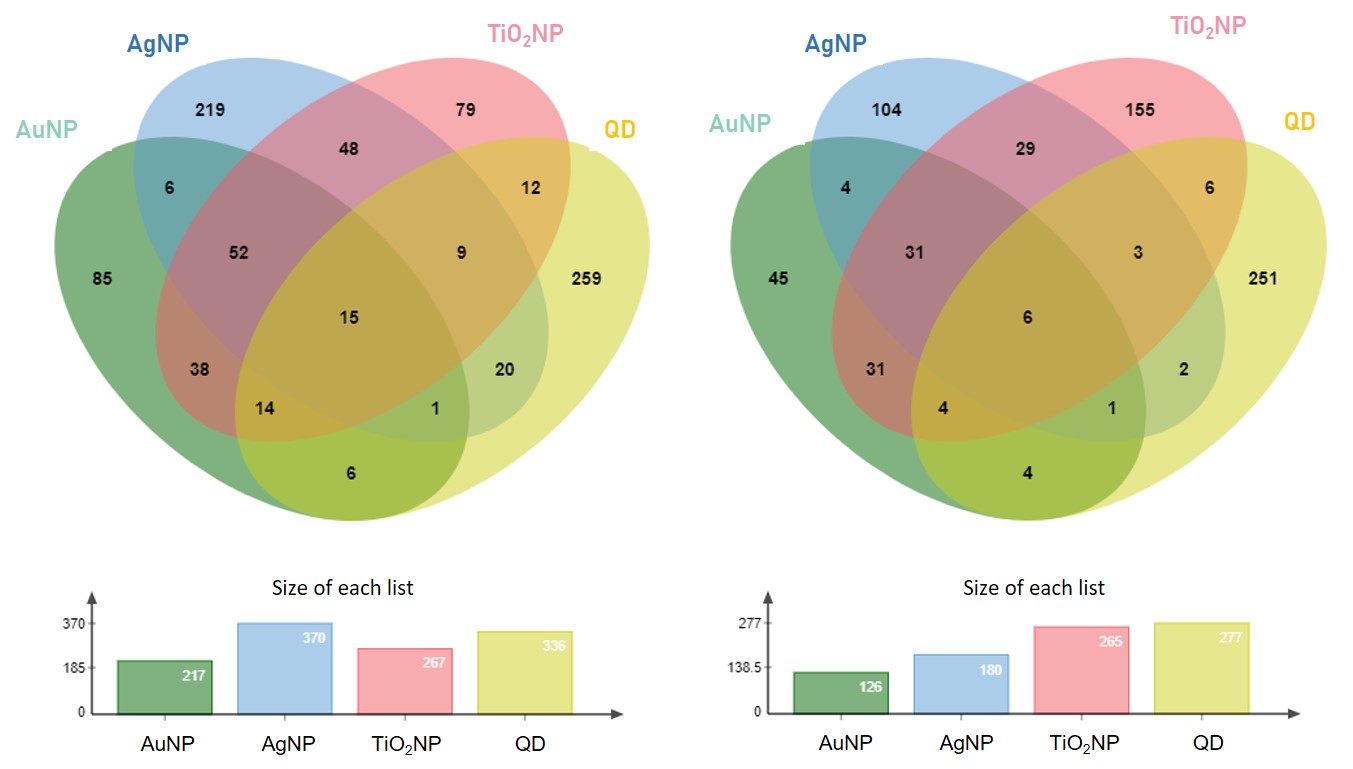

**Supplementary Figure 1**: Venn diagram of the common genes for all nanoparticles that are A) upregulated and B) downregulated.

|  | -log (p value) |
| --- | --- |
| Cellular Compartment nuclear speck | 1.01323 |
| Cellular Compartment mitochondrion | 1.01773 |
| Cellular Compartment extracellular space | 1.07572 |
| Cellular Compartment chloride channel complex | 1.20761 |
| Cellular Compartment perinuclear region of cytoplasm | 1.23657 |
| Cellular Compartment cytosol | 1.284 |
| Cellular Compartment extracellular exosome | 1.34679 |
| Cellular Compartment high-density lipoprotein particle | 1.49485 |
| Cellular Compartment Golgi apparatus | 1.95861 |
| Molecular Function collagen binding | 1.07058 |
| Molecular Function intracellular calcium activated chloride channel activity | 1.63827 |
| Molecular Function integrin binding | 1.74473 |
| Molecular Function peptidase activator activity | 1.79588 |
| Molecular Function protein binding | 1.82391 |
| Molecular Function chloride channel activity | 2.4318 |
| Biological Process muscle contraction | 1.02228 |
| Biological Process neuron apoptotic process | 1.07058 |
| Biological Process chloride transport | 1.08092 |
| Biological Process neuromuscular process controlling balance | 1.27572 |
| Biological Process positive regulation of monocyte chemotaxis | 1.58503 |
| Biological Process lipoprotein metabolic process | 1.61979 |
| Biological Process cell proliferation | 1.72125 |
| Biological Process negative regulation of cell proliferation | 1.85387 |
| Biological Process chloride transmembrane transport | 2.20761 |
| Biological Process cholesterol metabolic process | 2.40894 |

**Supplementary Figure 2**: Gene ontology analysis of common DEGs (p<0.05 and FC>2).

**Supplementary Table 2:** Genes that are related to the enriched GO terms for the common DEGs

| **GO Term** | **Gene Annotation** | **Related Genes** |
| --- | --- | --- |
| GO:0008203 | cholesterol metabolic process | APP, APOL1, CYP27A1 |
| GO:1902476 | chloride transmembrane transport | ANO6, APOL1, TTYH3 |
| GO:0008285 | negative regulation of cell proliferation | NDRG1, APP, ASPH, IGFBP6 |
| GO:0008283 | cell proliferation | ASPH, CERT1, TGFBI |
| GO:0042157 | lipoprotein metabolic process | APP, APOL1 |
| GO:0090026 | positive regulation of monocyte chemotaxis | APP, ANO6 |
| GO:0050885 | neuromuscular process controlling balance | APP, TPP1 |
| GO:0006821 | chloride transport | ANO6, TTYH3 |
| GO:0051402 | neuron apoptotic process | APP, PIGT |
| GO:0006936 | muscle contraction | ASPH, CERT1 |
